# Brain rhythms shift and deploy attention

**DOI:** 10.1101/795567

**Authors:** Craig G. Richter, Conrado A. Bosman, Julien Vezoli, Jan-Mathijs Schoffelen, Pascal Fries

**Affiliations:** Ernst Strüngmann Institute (ESI) for Neuroscience in Cooperation with Max Planck Society, Deutschordenstraße 46, 60528 Frankfurt, Germany; BCBL. Basque Center on Cognition, Brain and Language, Mikeletegi Pasealekua 69, 20009 Donostia, Spain; Donders Institute for Brain, Cognition and Behaviour, Kapittelweg 29, Radboud University, 6525 EN Nijmegen, Netherlands; Swammerdam Institute for Life Sciences, Center for Neuroscience, Faculty of Science, University of Amsterdam, Sciencepark 904, 1098 XH Amsterdam, Netherlands

**Keywords:** brain rhythms, theta, beta, gamma, attention, visual, cortex, electrocorticography

## Abstract

One of the most central cognitive functions is attention. Its neuronal underpinnings have primarily been studied during conditions of sustained attention. Much less is known about the neuronal dynamics underlying the processes of shifting attention in space, as compared to maintaining it on one stimulus, and of deploying it to a particular stimulus. Here, we use ECoG to investigate four rhythms across large parts of the left hemisphere of two macaque monkeys during a task that allows investigation of deployment and shifting. Shifting involved a strong transient enhancement of power in a 2-7 Hz theta band in frontal, pre-motor and visual areas, and reductions of power in an 11-20 Hz beta band in a fronto-centro-parietal network and in a 29-36 Hz high-beta band in premotor cortex. Deployment of attention to the contralateral hemifield involved an enhancement of beta power in parietal areas, a concomitant reduction of high-beta power in pre-motor areas and an enhancement of power in a 60-76 Hz gamma band in extra-striate cortex. Effects due to shifting occurred earlier than effects due to deployment. These results demonstrate that the four investigated rhythms are involved in attentional allocation, with striking differences between shifting and deployment between different brain areas.

**Significance:** We are often confronted by many visual stimuli, and attentional mechanisms select one stimulus for in-depth processing. This involves that attention is shifted between stimuli and deployed to one stimulus at a time. Prior studies have revealed that these processes are subserved by several brain rhythms. Therefore, we recorded brain activity in macaque monkeys with many electrodes distributed over large parts of their left hemisphere, while they performed a task that involved shifting and deploying attention. We found four dominant rhythms: theta (2-7 Hz), beta (11-20 Hz), high-beta (29-36 Hz) and gamma (60-76 Hz). Attentional shifting and deployment involved dynamic modulations in the strength of those rhythms with high specificity in space and time.

## Introduction

Selective attention is a central cognitive process that has been extensively studied both behaviorally and using invasive and non-invasive neurophysiological techniques. Despite widespread investigation, the mechanisms that control the shifting and deployment of attention are not yet fully understood. This is in part due to the fact that most respective studies compare attention conditions with no regard to temporal change, assuming static attention to different stimuli. Such approaches are blind to the temporally dynamic processes that construct an attentional state. Here, we investigate those dynamic processes, distinguishing between attentional shifting and attentional deployment. We define attentional shifting as the process that shifts attention in general, irrespective of the target stimulus or the shift direction; we define attentional deployment as the process that allocates attention to one out of two simultaneously present but spatially separate stimuli. There are a few studies that have investigated these questions non-invasively in humans. Among them, one study combined fMRI with an intricate task design to dissociate between brain activity underlying attentional deployment to a particular hemifield from activity underlying attentional shifts in general (1). The combination of event related fMRI and EEG with a specially designed attentional cueing paradigm allowed another study to track event-related potentials (ERPs) specifically related to attentional control or attentional orienting (2). Another study used steady-state evoked potentials to track attentional allocation in time and directly link the time courses of cortical facilitation to the behavioral benefits of attention (3). The current investigation seeks to build on these previous works by using high-resolution micro-electrocorticography (ECoG) in macaque monkeys to track the processes underlying attentional shifting and attentional deployment in space, time and frequency. The ECoG grid employed in this study provides an excellent means of tracking attentional dynamics, because it provides high spatial and temporal resolution while covering a large portion of sensory and executive cortical areas.

The need for high temporal resolution is underscored by recent studies showing that attention is subserved by brain rhythms for the control and implementation of attentional selection. Numerous studies have established that neurons across the hierarchy of visual areas show enhanced local and interareal gamma- and/or beta-band synchronization (4–17), and reduced theta-band synchronization and theta-gamma coupling (18), when processing attended as compared to un-attended stimuli. Yet, none of these studies has isolated the time-courses of attention-related effects from the time-course of sensory cue processing, or distinguished the time-course of general attentional shifting from the time-course of spatially specific attentional deployment. We hypothesize that attentional shifting and deployment are subserved by spatially, spectrally and temporally specific neuronal engagement. The current study employs a subtractive paradigm to isolate neurophysiological correlates of deployment and shifting of attention while minimizing cue-evoked confounds. Our results show a remarkable diversity of function across four distinct narrow-band rhythms in the theta, beta, high-beta and gamma bands. This activity shows distinct temporal dynamics across frequencies that arise in specific cortical locations.

## Results

### Two contrasts, isolating attentional deployment and attentional shifting

Two macaque monkeys performed a task entailing spatially selective visual attention (17), which is illustrated and described in detail in Figure 1A. Electrocorticographic (ECoG) grid recordings were obtained from a large portion of the left hemisphere (Fig. 1B). Two stimuli were presented in the two visual hemifields, and at a later time, one of them was cued to be the behaviorally relevant target, leaving the other one to be the distracter. Monkeys were required to report with a bar release randomly timed changes in the target. The task paradigm allowed for changes in either stimulus also prior to cue onset, and in this case, responses were rewarded in a random 50% of the trials.

**Fig. 1.**
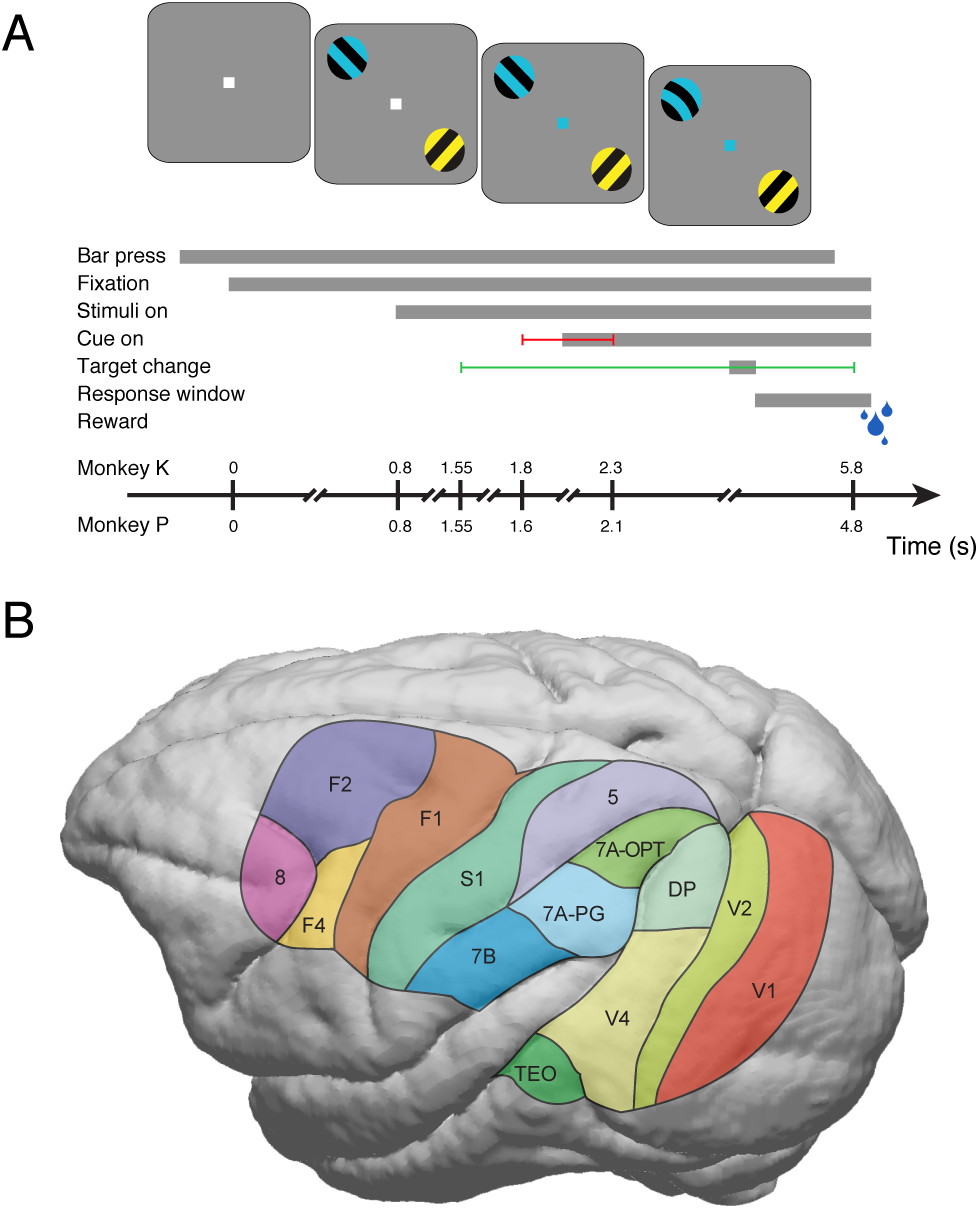
Behavioral task and regions of interest. (*A*) Schematic of a correct trial with attention directed to the blue stimulus in the hemifield ipsilateral to the recording grid. Trials commenced with the monkey touching a bar. This triggered the presentation of a fixation point. Monkeys were required to maintain their gaze within a prescribed fixation window throughout task performance (monkey K: 0.85 deg radius, monkey P: 1 deg radius); otherwise the trial terminated and a timeout was given before the next trial started. Following a fixed interval (intervals shown as a timeline at the bottom), two isoluminant and isoeccentric drifting sinusoidal gratings appeared, one in each hemifield (diameter: 3 deg, spatial frequency: ≈1 cycle/deg, drift velocity: ≈1 deg/s, resulting temporal frequency: ≈1 cycle/s, contrast: 100%). Blue and yellow tints were randomly assigned to each of the gratings on each trial. Following a variable duration (indicated by horizontal red line), the fixation point changed color to match one of the stimuli, indicating the stimulus to be covertly attended, which we refer to as the target. The un-cued, behaviorally irrelevant stimulus is referred to as the distracter. Either one of the two stimuli could undergo a transient change (bending of grating stripes as illustrated), lasting 150 ms. This change could occur within the longer time period indicated by the horizontal green line. For each trial, two time points were drawn from a slowly increasing Hazard rate, and randomly assigned to the target and distracter change. As a consequence, the first stimulus change in a trial was equally likely to be a target or a distracter change, and these changes occurred with identical temporal probabilities. If the distracter change occurred before the target change, monkeys were required to wait until the target change and report it with a bar release. Stimulus changes could occur both before and after the cue. Stimulus changes before the cue were included to capture the time course of the attentional deployment after cue onset. Once the cue had been given, bar releases in response to changes of the target were rewarded. Before the cue, bar releases in response to changes of either stimulus were rewarded in a random 50% of the cases. The response window was 150-500 ms after the start of the respective stimulus change. (*B*) The joint coverage over the two monkeys of the selected brain areas. The figure shows the coverage obtained in both monkeys; regions covered in only one monkey are excluded. Site coverage has been co-registered to a common macaque template brain.

Unexpectedly, the monkeys showed a spontaneous bias towards responding to blue stimuli prior to cue onset (Fig. 2A; χ^2^(1,N=793) = 88.59, p = 0). This bias disappeared after cue presentation, when the monkeys showed a balanced response profile to blue and yellow target stimuli (Fig. 2B; χ^2^(1,N=2296) = 2.31, p = 0.13). This bias was also reflected in the reaction times of the monkeys, where responses to yellow targets were significantly longer than those to blue targets up to 150 ms after cue onset (Fig. 2C). Taken together, the greater likelihood of reporting pre-cue changes in blue rather than yellow stimuli, and the longer reaction times to post-cue changes in yellow targets indicate the presence of a spontaneous attentional bias toward the blue stimulus prior to cue onset.

**Fig. 2.**
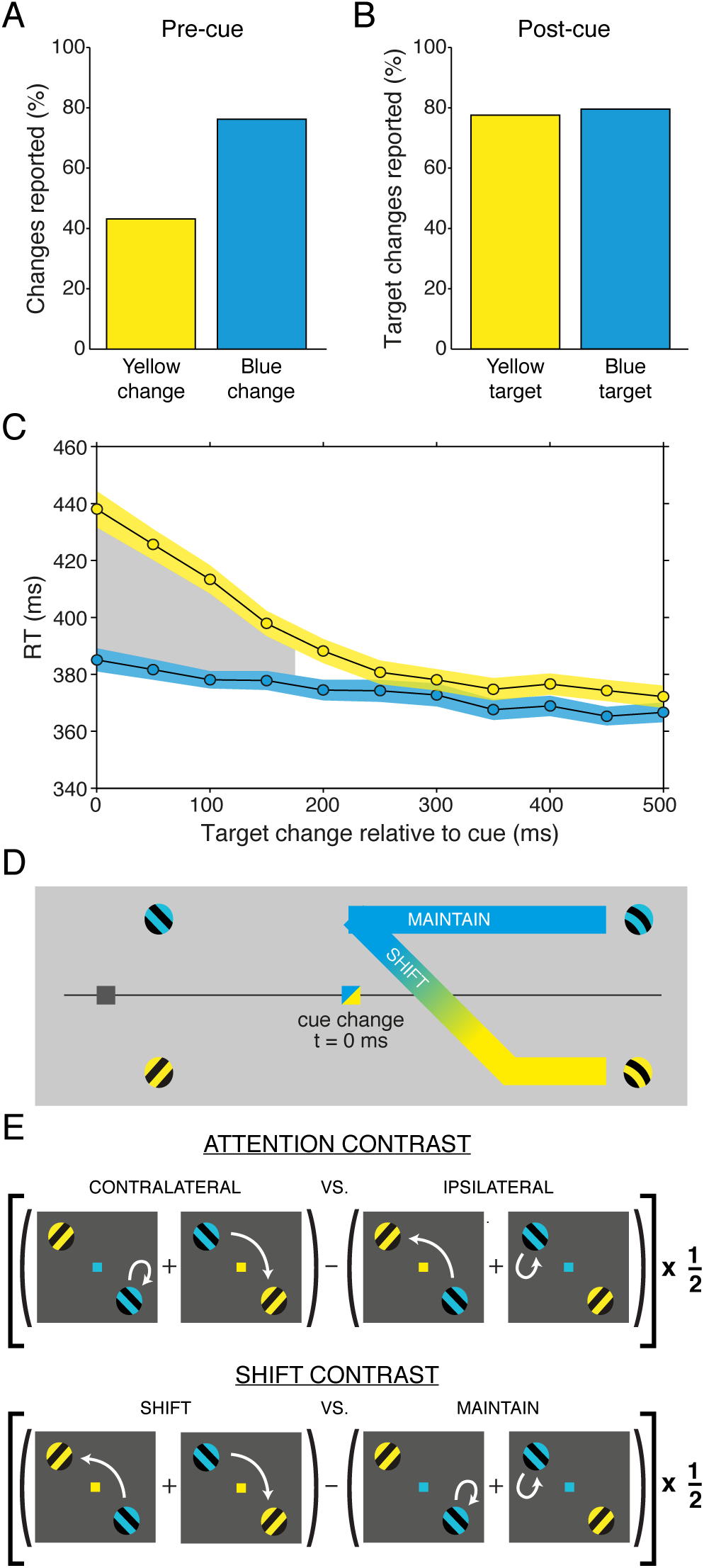
Behavioral analysis and condition contrasts used for the neurophysiological analysis. (*A*) The behavioral response pattern before cue presentation showed a bias towards responses to the blue stimulus (chi-squared test: χ^2^(1,N=793) = 88.59, p = 0). (*B*) Following the presentation of the attentional cue, the monkeys showed no significant difference in response rates to blue or yellow target stimuli (χ^2^(1,N=2296) = 2.31, p = 0.13). (*C*) Reaction times as a function of the latency between cue presentation and the start of the target change for the yellow (yellow line) and blue (blue line) stimuli. Reactions times were binned over 200 ms regions, at 50 ms intervals. Colored shaded regions indicate ±1 SEM, pooled over the two monkeys. The gray-shaded region indicates a significant difference in reaction time to blue versus yellow stimuli (p<0.05, two-tailed non-parametric randomization test, corrected for multiple comparisons across time windows). (*D*) Schematic diagram of the monkeys’ inferred attentional location as a function of time around cue onset. Cue onset signals that the monkey must change the favored behavioral response (switch) or maintain the current bias (stay). (*E*) Each gray square illustrates one of the four possible combinations of stimulus and fixation point coloring. Data from these task conditions were combined as illustrated by the mathematical formula made up of the individual conditions. This resulted in two contrasts, the **attention contrast** and the **shift contrast**, as explained in detail in the results section. Each contrast was multiplied by a factor of 1/2 to preserve the original magnitudes of the measurements.

Figure 2D illustrates the inferred attentional location as a function of time around cue onset. This depicts the attentional bias to the blue stimulus until cue onset, at which point a blue cue indicates that the monkey must maintain attention to the blue stimulus, whereas a yellow cue indicates that attention should be shifted to the yellow stimulus. We took advantage of this unexpected, spontaneous bias and constructed two contrasts for the analysis of the ECoG recordings, as illustrated in Figure 2E. One contrast is referred to as the **attention contrast**, because it isolates the effects of deploying selective attention to the contralateral versus the ipsilateral hemifield. Effects of attentional shifting as such are contained in the individual component conditions but are removed by the subtraction. The other contrast is referred to as the **shift contrast**, because it contrasts the shifting of attention in either direction with maintaining attention in either hemifield, and thereby isolates the effects of attentional shifting. Effects of the deployment or maintenance of attention to a particular hemifield are contained in the individual component conditions, but are removed by the subtraction.

Note that the **attention contrast** is completely balanced with regard to stimulus and fixation point coloring. That is, each stimulus is colored both blue and yellow on each side of the subtraction. Similarly, the fixation point is colored both blue and yellow on each side. This ensures that any differences in the neuronal representation of stimulus or fixation point color are not mistaken as attention effects.

Note that the **shift contrast** is not completely balanced. Balancing is achieved for stimulus coloring, but not for fixation point coloring. We acknowledge this fact here, while in the discussion section, we explore the likelihood of this imbalance to explain the results obtained with the **shift contrast**.

### Oscillatory activity is most prominent in four frequency bands

Oscillatory activity can be detected with particularly high sensitivity by metrics of phase coupling (19, 20). Therefore, the pairwise phase consistency (PPC) (21) was computed between all possible pairs of recording sites, and the peaks in each PPC spectrum were identified via an automated fitting algorithm (Fig. 3B,D). This showed that in both monkeys, the probability of finding a peak was highest within four distinct frequency bands (Fig. 3A,C). These bands corresponded to the previously described theta, beta, high-beta and gamma rhythms. Each band was characterized by its peak frequency and the full-width-at-half-maximum (see Fig. 3A,C), which defined four frequency bands of interest (FOIs). Analyses focused on local field potential (LFP) power, averaged within those FOIs and over monkeys. We investigated the temporal dynamics for the **attention contrast** and the **shift contrast**, separately for the four FOIs and the 14 brain areas illustrated in Figure 1B.

**Fig. 3.**
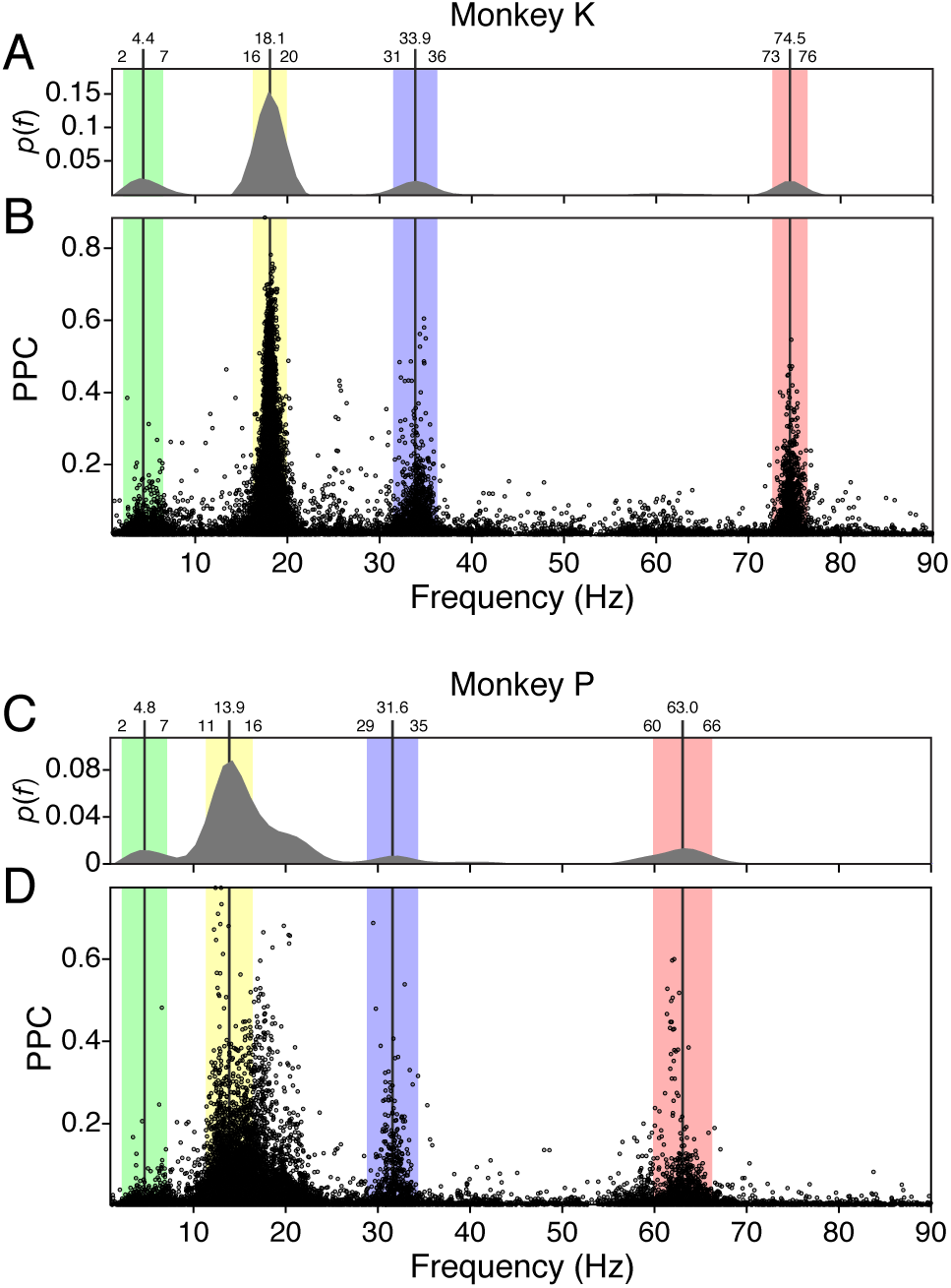
Determination of individual spectral peaks per monkey. (*B*) Each dot corresponds to a peak found in the spectrum of phase locking (PPC) between a given site pair of monkey K. Each spectrum could contain multiple peaks. There were 21115 site pairs in this monkey. (*A*) probability mass of detecting PPC peaks as a function of frequency in monkey K. Black vertical lines and the corresponding frequencies noted at the top indicate estimated peaks of the probability distribution. Colored regions and the corresponding frequencies noted at the top denote the full-width-at-half-maximum for each detected peak. The peaks and the full-width-at-half-maximum defined the frequency bands of interest (FOIs). (C,*D*) Same conventions as (A,*B*), for monkey P (20503 site pairs). Both monkeys showed distinct regions of theta (green), beta (yellow), high-beta (blue), and gamma (red) frequency oscillatory activity.

### Attention contrast and shift contrast in time and space: theta

Figure 4B shows the dynamics of theta power in the 14 areas, separately for attentional deployment to the stimulus contralateral (red) and ipsilateral (green) to the recorded hemisphere. The respective statistical testing entailed correction for the multiple comparisons across FOIs, areas and the investigated time points. This contrast did not reveal significant differences. Note that a previous study established an attentional reduction in theta activity within V1 and V4, theta synchronization between V1 and V4 and theta-gamma coupling in V1(18). This study primarily used data from later periods relative to cue onset. Consistent with that, our present analyses show the same trend for an attentional reduction in theta power in V1, V2, V4 and TEO, towards the end of the present analysis window. Figure 4A illustrates the respective topographical distributions. Here and in the following, these topographical plots are provided for illustration of the full spatial distribution of power contrasts without statistical testing, whereas statistical testing is provided for the time-courses of the separate areas.

**Fig. 4.**
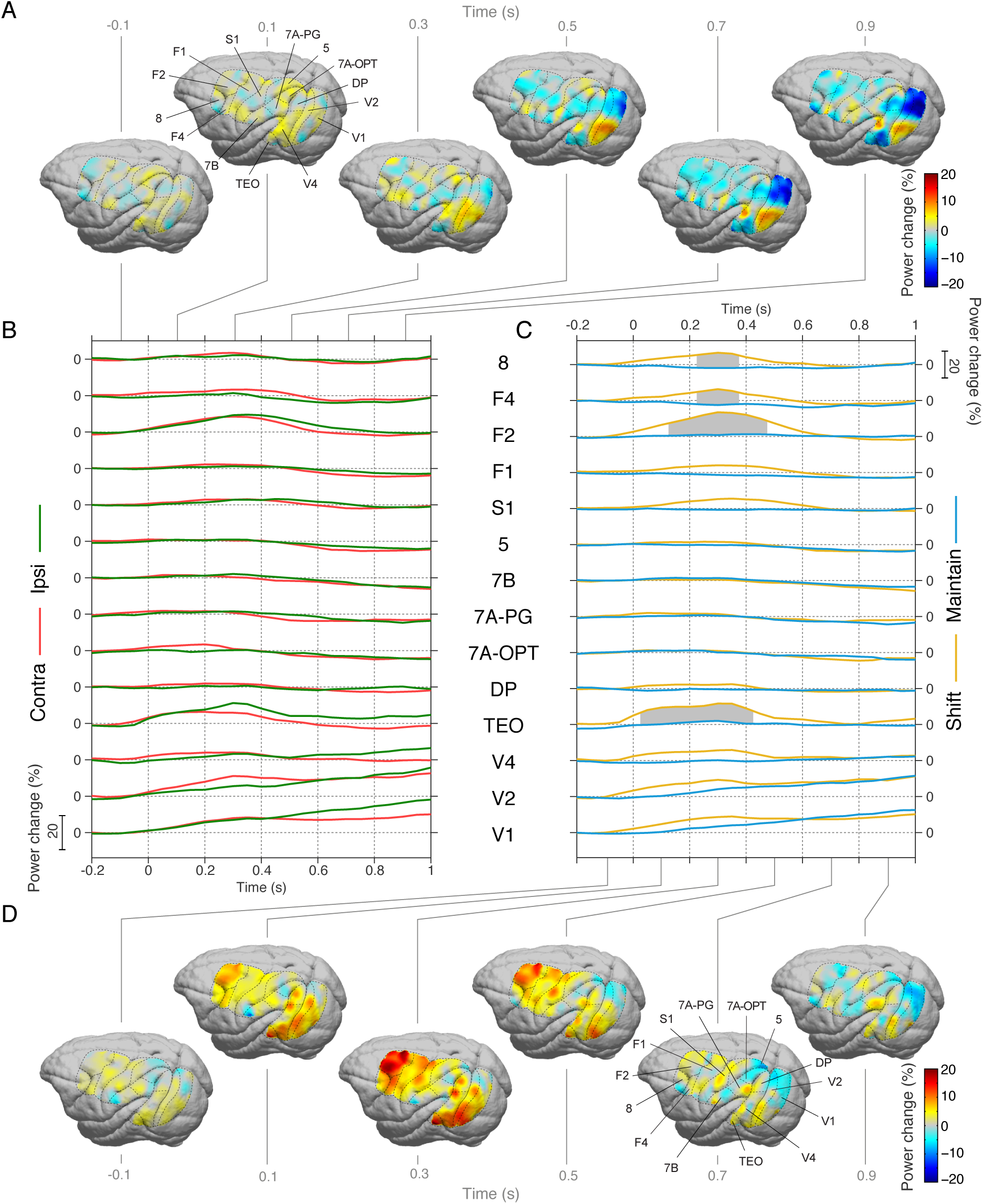
Attention and shift contrasts in time and space: theta. (*A*) Topography of the **attention contrast** for theta-band power, for 200 ms windows centered on the indicated times, averaging both monkeys on a common macaque template (un-thresholded). Brain area delineations are marked by dashed lines. (*B*) Percentage power change from baseline (200 ms window prior to cue onset) for attentional deployment to the contralateral (red line) and ipsilateral (green line) stimulus, averaged over all sites within each of the indicated brain areas and then over monkeys. For each brain area, the gray dotted line indicates zero change relative to baseline. For all brain areas jointly, the scale indicates the magnitude of the power as a percentage change from baseline. Significant differences between conditions are denoted by gray shading (p<0.05, two-tailed non-parametric randomization test, corrected for multiple comparisons across time windows and brain areas, and Bonferroni corrected for the four frequency bins). (*C*) Same as *B,* but showing percentage change from baseline for trials in which attention was shifted (yellow line) or maintained (blue line). (*D*) Same as *A*, but for the **shift contrast**.

Figure 4C shows the corresponding analyses for the shifting (yellow) versus maintaining (blue) of attention. **Shifting** induced a transient enhancement of theta power peaking between 200 and 400 ms after cue onset, in the pre-frontal and pre-motor areas. A very similar enhancement was significant in TEO and trending across visual areas, even though these areas are distant to the frontal areas with regard to their spatial position and distinct with regard to their beta dynamics (see below). This transient theta power enhancement did not reach significance in sensorimotor areas F1 and S1 and in area DP, even though these areas are partly neighboring the frontal areas and exhibiting similar beta dynamics (see below). When attention was maintained, theta power dynamics lacked any notable transient change. Figure 4D illustrates the respective topographies.

### Attention contrast and shift contrast in time and space: beta

Figure 5B shows the dynamics of beta-band power for the **attention contrast**. Attentional deployment to the contralateral stimulus induced an enhancement of beta-band power in posterior parietal areas DP and 7A-OPT, reaching significance at approximately 650 ms, and persisting to the end of the analysis window at 1 s. Frontal and pre-motor areas F2, F4 and area 8 showed a similar trend. Figure 5A shows the respective topographies.

**Fig. 5.**
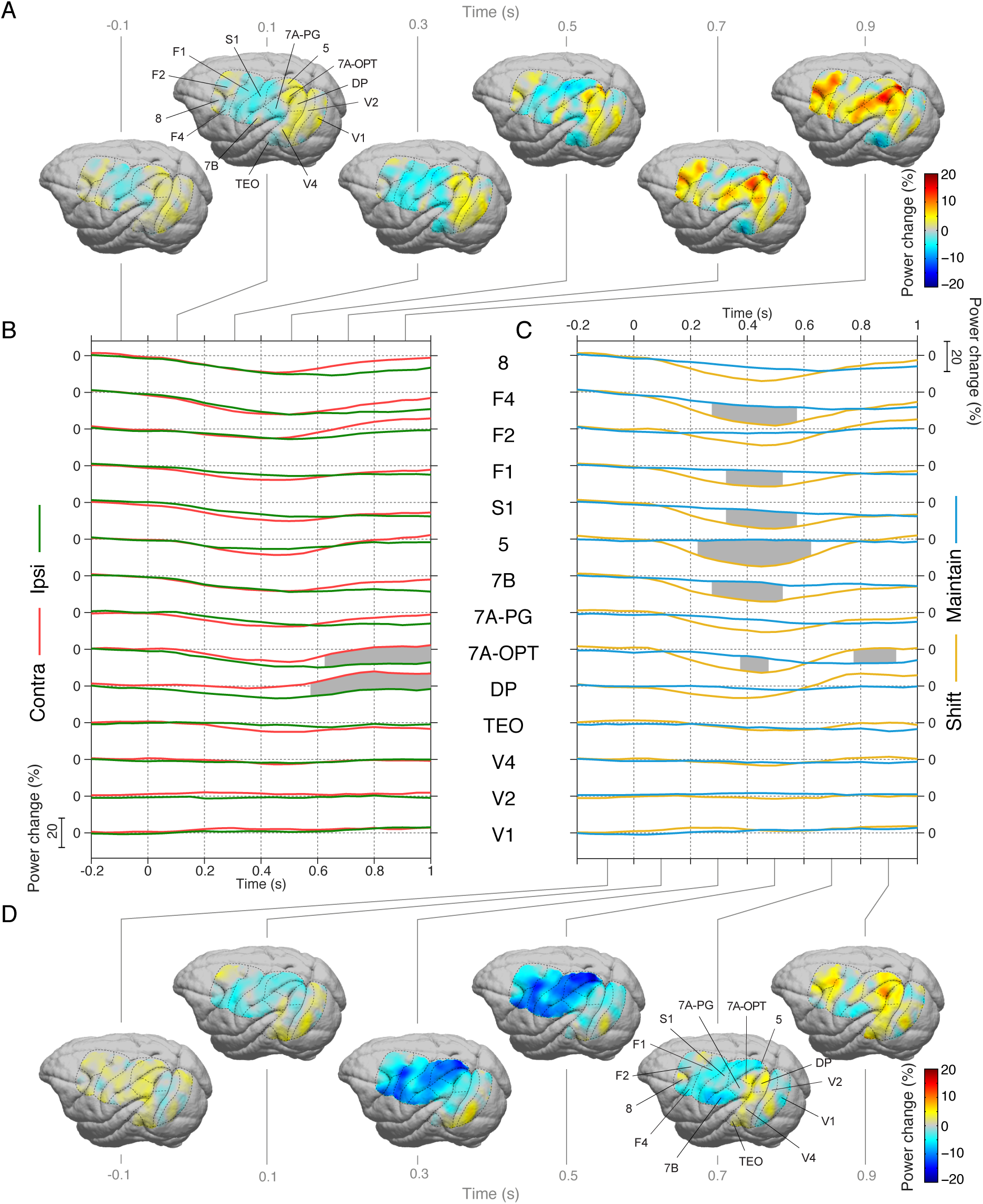
Attention and shift contrasts in time and space: beta. Same conventions as Fig. 4.

Figure 5C shows the beta-band dynamics for the **shift contrast**. Attentional shifting is associated with a transient pronounced decrease in beta frequency power, peaking at approximately 400 ms post-cue, which is highly consistent across frontal, pre-motor, sensorimotor and posterior parietal areas. When attention was maintained, beta power dynamics lacked any notable transient change, similar to the above described theta-band power dynamics. Yet note that the signs of the transient, short-latency shifting effects were opposite, with beta power decreases and theta power increases. Figure 5D shows the corresponding topographies for the **shift** dynamics.

### Attention contrast and shift contrast in time and space: high-beta

Figure 6B shows the dynamics of high-beta-band power for the **attention contrast**. Attentional deployment to the ipsilateral stimulus, i.e. an attentional disengagement of the recorded hemisphere, induced an increase in high-beta power in areas F2, F4 and TEO. Note that while pre-motor areas showed a clear high-beta peak in their power spectra, this was not the case for TEO. Note also that attention increased high-beta power in some areas (Fig. 6B) while decreasing beta power in others (Fig. 5B). The topographies for the high-beta **attention contrast** are shown in Figure 6A.

**Fig. 6.**
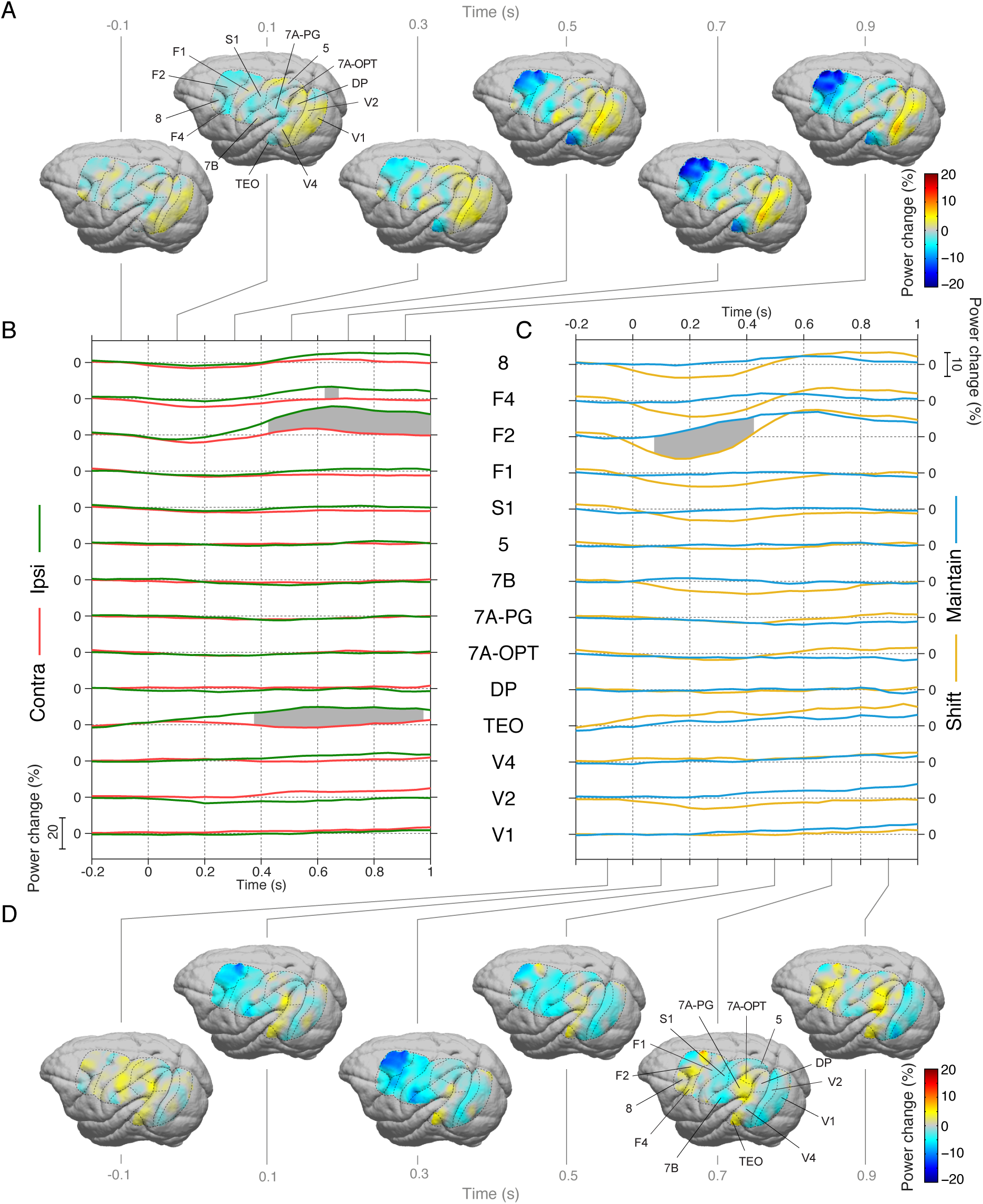
Attention and shift contrasts in time and space: high-beta. Same conventions as Fig. 4,5.

Figure 6C shows the high-beta power dynamics associated with the **shift contrast**. Attentional shifting induced a transient decrease in high-beta power in area F2, peaking at ≈250 ms. Similar trends are present in areas F4, area 8, F1 and S1. This effect is similar to that seen for beta power in neighboring areas (Fig. 5C), though the high-beta power decrease is maximal approximately 150 ms earlier and is spatially more constrained. Figure 6D illustrates the topographical evolution of the dynamics for the **shift contrast**.

### Attention contrast and shift contrast in time and space: gamma

Figure 7B depicts the time course of gamma power differences for the **attention contrast**. Attentional deployment to the contralateral stimulus induced enhanced gamma power in extrastriate areas. Latencies of this enhancement were shorter for areas higher up in the visual hierarchy, i.e. there was a backward progression of attentional effects as shown before for firing rates (22): After cue presentation, enhancements reached significance at 450 ms in TEO, 500 ms in V4 and 600 ms in V2. Unlike in the beta and high-beta bands, attentional effects in the gamma band were confined to extrastriate cortex and were not detectable in more anterior areas. The associated power change topographies are shown in figure 7A.

**Fig. 7.**
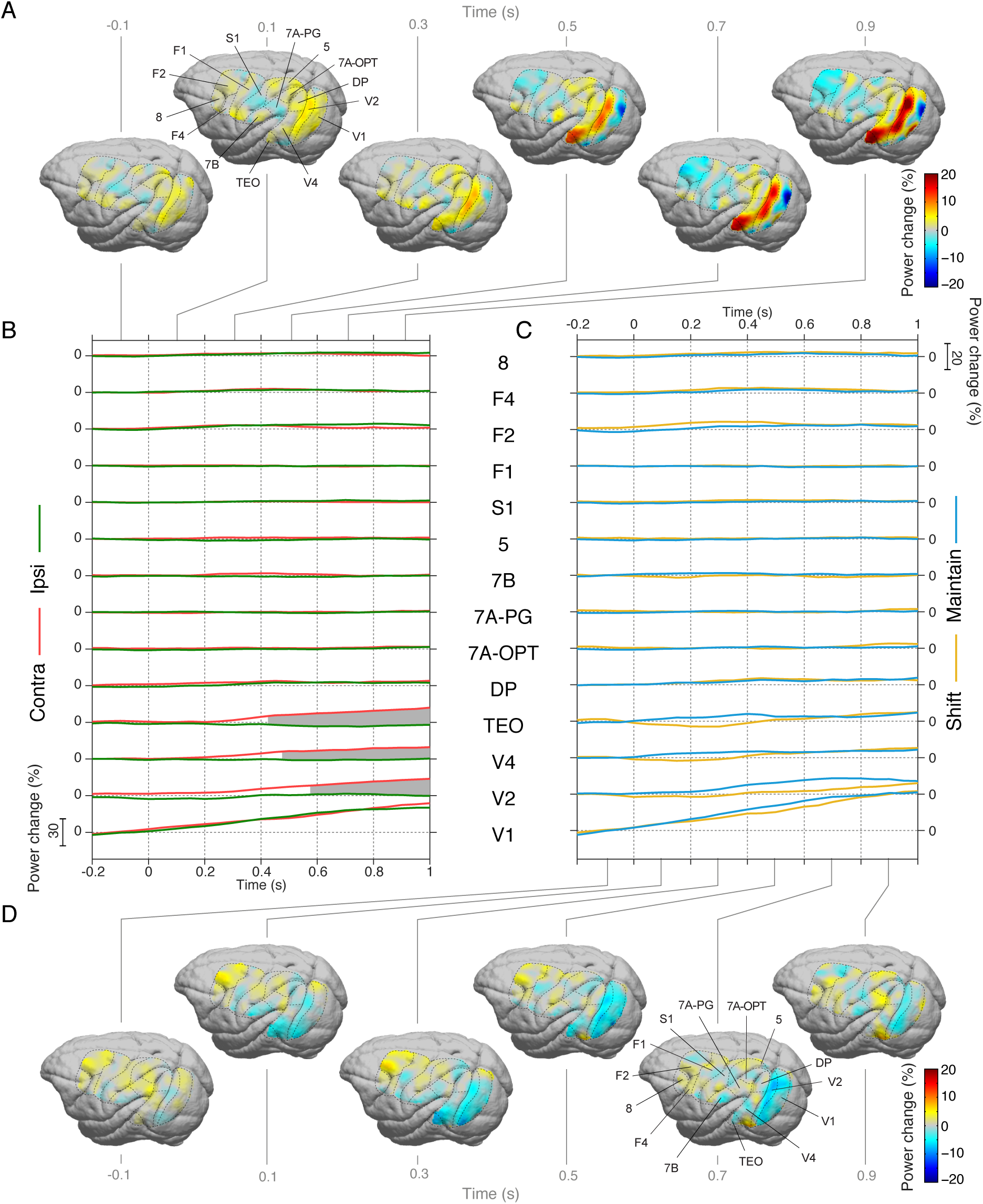
Attention and shift contrasts in time and space: gamma. Same conventions as Fig. 4,5,6.

Figure 7C shows the **shift contrast** for gamma power and reveals no significant differences. The associated power difference topographies are shown in figure 7D.

### Shift effects occurred earlier than deployment effects

A closer inspection of the time-courses of effects suggested that overall, the differences due to attentional shifts occurred earlier than the differences due to attentional deployment. To test this, we compiled a metric of overall differences separately for the shift and the deployment contrast: We rectified the condition differences, averaged them over areas and frequency bands and tested whether this value was significantly larger than zero (non-parametric randomization of conditions across trials, corrected for the multiple comparisons over time points). Figure 8 shows the resulting time courses and confirms that the overall shift effect starts earlier than the overall deployment effect. The shift effect reaches significance at the time of the cue presentation. This is possible, because each indicated time point corresponds to an analysis window of ±250 ms length. Furthermore, the shift effect shows a peak around 300 ms after the cue. The deployment effect reaches significance at 200 ms after cue presentation, and it steadily increases with time after the cue.

**Fig. 8.**
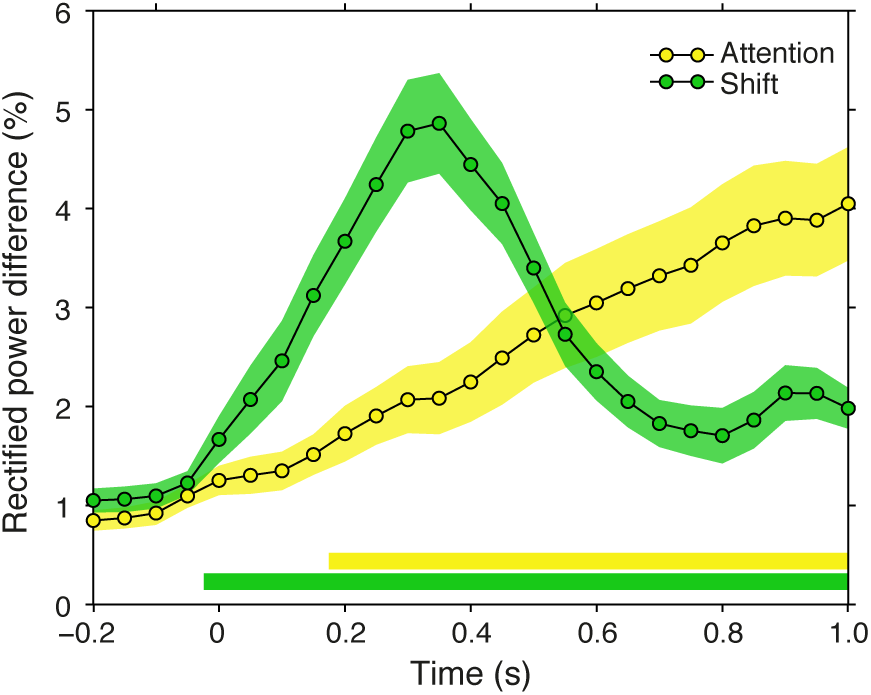
Shift effects occur earlier than deployment effects. The rectified power difference averaged over frequency bands, brain areas and monkeys shown for the **shift contrast** (green), and the **attention contrast** (yellow). Green and yellow horizontal bars on the bottom denote the period that the rectified difference is statistically significant for the shift and attention contrasts, respectively (p<0.05, two-tailed non-parametric randomization test, corrected for multiple comparisons across time windows). Colored shaded regions indicate ±1 SEM computed across brain areas.

## Discussion

In summary, we used large-scale high-density ECoG in two macaque monkeys and analyzed the signals, differentiating them in space, time and frequency, to test for effects of attentional deployment or shifting. This revealed four rhythms that showed effects with clear spatial, temporal and spectral specificity.

The spatial specificity was reflected in the fact that different frequency bands showed very different effects in different brain areas: Theta effects, which were exclusively significant for shift contrasts, occurred in areas 8, F4, F2 and TEO, while sparing high-level areas in parietal cortex, like areas 7A-OPT, 7A-PG and 7B; high-beta power effects were primarily localized to F2 and F4; gamma effects were restricted to extrastriate visual areas TEO, V4 and V2.

The temporal specificity was apparent in the fact that shift effects occurred earlier than deployment effects. Among shift effects, both theta and high-beta effects tended to occur earlier than beta effects.

The spectral specificity was evident in the fact that beta and high-beta showed shift effects in the same direction, yet deployment effects in the opposite direction. Also, there were opposite shift effects for theta versus beta and high-beta. While theta showed a shift-related enhancement, beta and high-beta showed a shift-related decrease. This latter observation supports the notion that beta is involved in the maintenance of the status-quo (23) and is therefore reduced when attention shifts; it might also support the notion that theta is involved in shifting in the sense of an attentional reset (24, 25).

Essentially the only case, in which two rhythms showed a similar effect is the shift-related reduction in beta in area F4 and of high-beta in area F2; yet even there, the effects began earlier in high-beta than beta; furthermore, for the deployment contrast, the same rhythms in the same areas showed opposite effects or trends. Thus, our observation that the effects differ at least along space or time, or between the shift and deployment contrast, strongly suggests that the different rhythms are regulated by independent mechanisms. Note that studies relying solely on conventional metrics of neuronal activation, like neuronal firing rates or BOLD, would not be able to see the differential and sometimes opposing effects on different rhythms, and the concomitant spatial and temporal specificity of those effects. This demonstrates the usefulness of large-scale high-density ECoG recordings, allowing analyses that are resolved simultaneously along the spatial, temporal and spectral dimension.

A point of potential concern relates to the imbalance of the cue properties for the shift contrast. Unlike the attention contrast, which is fully balanced in stimulus and cue properties, the shift contrast has unbalanced cue colors, such that attentional shifting occurs in response to the yellow fixation point, while the maintenance of attention is triggered by the blue fixation point (Fig. 2E). We argue that this imbalance is unlikely to explain the majority of the observed effects. The physical difference between a yellow versus a blue fixation point is expected to cause local effects in neurons that are selective for the representation of the fovea and that are color selective. Neurons selective for different colors are partly intermingled within cortical areas (26, 27), such that our recordings with 1 mm diameter ECoG electrodes might well average over different color domains and thereby reduce or even eliminate color-differential responses. Potential residual color-differential responses in individual recording sites should be further reduced by our averaging over all recording sites in a given area. In contrast to those expectations for color-differential responses, the cognitive difference between a yellow versus a blue fixation point, i.e. the attentional shift, is expected to cause widespread effects, including pre-frontal and pre-motor areas. Our results are consistent with this expectation, because we find near-simultaneous effects in those areas and visual areas. The behavioral data suggest that both animals, in the period before cue onset, spontaneously chose to attend the blue stimulus, probably because it was more salient and/or it allowed an easier detection of the to-be-reported shape change. If the same preference for blue would have led to stronger responses to the fixation point turning blue, then this should have induced stronger spectral perturbations in response to blue cue onsets. By contrast, we find that spectral perturbations were stronger for yellow cue onsets. This is particularly prominent in the theta and beta bands, where blue cues hardly or not at all perturbed the dynamics, whereas yellow cues led to very clear transient perturbations lasting for 0.2-0.4 s. This pattern suggests that the effects of the two cue colors are to be interpreted in the shift-versus-maintain sense, because maintaining attention (blue) is expected to involve less cognitive effort than shifting it (yellow). Only for gamma in striate and extrastriate areas did we find a trend towards enhancement with the blue cue, as expected for an effect of higher salience, yet this did not reach significance.

Several previous investigations have explored some aspects related to the present study. The time course of attentional shifts has been investigated with steady-state visual evoked potentials (SSVEPs) obtained with EEG recordings from human subjects performing an attention task similar to our task (3). In response to cue presentation, SSVEPs showed neuronal signs of attentional shifting with close temporal relationship to the attentional effect on behavior. The spatial pattern of brain regions involved in attentional deployment and shifting has been investigated with fMRI in human subjects (1). BOLD signals in extrastriate cortex reflected attentional deployment for the duration of sustained attention. By contrast, BOLD signals in posterior parietal cortex were transiently enhanced during attentional shifts. The BOLD signal is correlated to different measures of neuronal activation, and is particularly strongly related to gamma-band activity (28–31). In agreement with this and the fMRI study, we found gamma to be enhanced for the attention contrast in extrastriate cortex, starting around 0.4 s after cue onset and lasting until the end of the analysis period. Note that our analysis of beta and high-beta revealed additional effects of sustained attentional deployment outside of visual cortex, in areas DP and 7A-OPT for beta, and in areas F2 and F4 for high-beta. Note also that our analysis did not reveal a shift-related transient increase in posterior parietal gamma, as might have been expected on the basis of the fMRI results and other studies showing attention-related parietal gamma modulation (32). Our ECoG might not have covered the involved parts of parietal cortex, which might be located inside the intra-parietal sulcus, and/or it might have had too low spatial resolution to reveal very local gamma enhancements. Another study used fMRI to aid the analysis of event-related potentials (ERPs) during cue-related attentional deployment (2). This revealed an activation sequence starting in medial frontal cortex, progressing through medial parietal cortex and finally affecting visual occipital cortex. The relations between ERPs and long-lasting perturbations of different rhythms are not well understood. An analysis of ERPs in the current ECoG dataset will allow a more direct comparison with the human ERP data and is a highly relevant task for the future. A similar sequence of frontal-then-visual engagement as described with the ERPs has also been found with combined microelectrode recordings from the frontal eye field (FEF) and V4 in macaques (6). Attention enhances gamma Granger causality from FEF to V4 as soon as 110 ms after cue presentation, whereas it enhances gamma Granger causality from V4 to FEF only from 160 ms onwards. Also, studies investigating attention effects on firing rates have found similar sequences. Firing rate enhancements in response to attentional targets occur first in prefrontal and subsequently in parietal cortex (15). Similarly, firing rate enhancements in response to attended versus non-attended stimuli occur earlier in V4, at intermediate latencies in V2 and at the longest latencies in V1 (22).

Future work will need to investigate putative cross-frequency interactions between the rhythms described here (18). For example: Does the timing and strength of the shift-related pre-frontal and pre-motor theta enhancement on a given trial predict the timing and strength of the shift-related high-beta and beta decreases in those regions and/or the beta decreases in parietal areas? How are high-beta and beta related, given that they show partly similar and partly opposite dynamics, and that they occupy partly the same territory (pre-frontal and pre-motor), yet partly different territory (parietal shows beta effects, but no high-beta effects). The present and those future investigations have been made possible through the simultaneously high spatial and temporal resolution of the high-density large-scale ECoG approach. Yet, as mentioned above, further improvements in density will likely reveal further detail e.g. in parietal and pre-frontal cortex. As an isotropic increase in density will lead to a cubic increase in channel count, future approaches will likely have to find a compromise between coverage and density, and combine widespread low-density with targeted high-density recordings.

## Methods

### Paradigm, stimulation and subjects

Data from two adult male macaque monkeys (macaca mulatta) were collected for this study. All experimental procedures were approved by the ethics committee of the Radboud University Nijmegen (Nijmegen, The Netherlands). Stimuli were presented on a CRT monitor (120 Hz non-interlaced) in a dimly lit booth and controlled by CORTEX software (https://www.nimh.nih.gov/labs-at-nimh/research-areas/clinics-and-labs/ln/shn/software-projects.shtml). The paradigm with all details is illustrated in Figure 1A and its legend.

### Electrophysiological recording and preprocessing

LFP recordings were made via a 252 channel electrocorticographic grid (ECoG) subdurally implanted over the left hemisphere (33). Data from the same animals, overlapping partly with the data used here, have been used in several previous studies (7, 16, 17, 34–41). Recordings were sampled at approximately 32 kHz with a passband of 0.159 – 8000 Hz using a Neuralynx Digital Lynx system. The raw recordings were low-pass filtered at 250 Hz, and downsampled to 1 kHz. The electrodes were distributed over eight 32-channel headstages and referenced against a silver wire implanted onto the dura overlying the opposite hemisphere. The electrodes were re-referenced via a bipolar scheme to achieve **1**) greater signal localization 2) cancellation of the common reference, 3) rejection of headstage specific noise. The bipolar derivation scheme subtracted the recordings from neighboring electrodes (spaced 2.5 mm) that shared a headstage, resulting in 218 bipolar derivations, referred to as “sites” (see (16) for a detailed description of the re-referencing procedure).

All signal processing was conducted in MATLAB (MathWorks, USA) and made use of the FieldTrip toolbox (http://www.fieldtriptoolbox.org/) (42). Raw data were cleaned of line noise via the subtraction of 50, 100, and 150 Hz components fit to the data using a discrete Fourier transform. Trial epochs for each site were de-meaned by subtracting the mean over all time points in the epoch. Sites with excessive noise or lack of signal were excluded, leaving 207 of 218 sites for monkey K, and 203 of 218 for monkey P. Epochs with any site having a variance of greater than 5 times the variance based on all data from that same site in the same session were rejected. In addition, epochs were manually inspected and epochs with artifacts were rejected. Subsequently, all epochs were normalized such that the concatenation of all epochs for a given site had a standard deviation of 1. Following this, all epochs of each site were combined across sessions.

### Region of interest definition

Fourteen brain areas, shown in Figure 1B, were selected for analysis. Brain area definitions were defined as follows: 1) Each monkey’s electrode locations were aligned with its respective anatomical MRI, based on sulcal locations from high resolution intraoperative photographs. The MRI and electrode locations were then warped to the F99 template brain in CARET (43), such that each electrode location could be compared with anatomical atlases provided by the CARET software. Based on these atlases, bipolar derivations with both electrodes within the same area were assigned to that area (see (16) for a more detailed description).

Spatial maps have been restricted to show the average activity across monkeys only at those locations, where both monkeys had ECoG grid coverage after co-registration. Spatial maps are shown on the INIA19 macaque brain (44) after co-registration of this template and each monkey’s site locations to the F99 template brain in CARET (43).

### Segmentation of data into analysis periods

All analyses were computed on correctly performed trials, i.e. where a response was logged within the allotted time interval after the target change.

To identify the most prominent frequency bands, we used phase locking analysis employing the pairwise phase consistency (PPC) metric (21). For this analysis, the data from 300 ms after cue onset until a target or distracter change was segmented into 500 ms epochs with 60% overlap. The first 300 ms after cue onset were excluded to avoid transients. As target and distracter changes occurred at randomized times, this resulted in a variable number of epochs per trial. Overlap was employed to implement Welch’s method (45) for improving spectral estimation and optimized for use with the multitaper method (46, 47). This procedure resulted in 15518 epochs (monkey K: 6689, monkey P: 8829).

After identifying the most prominent frequency bands with PPC analysis, subsequent analyses focused on time-varying power in those bands. Time-varying power was analyzed for periods beginning 450 ms prior to cue onset and ending when a change occurred in either the target or distracter stimulus. As target and distracter changes occurred at randomized times, this resulted in periods of variable length. This resulted in 4722 epochs (monkey K: 2190, monkey P: 2532), with a mean length of 1669 ms and a standard deviation (SD) of 912 ms (monkey K: 1551 ms, SD = 903 ms, monkey P: 1771 ms, SD = 907 ms). Periods were approximately evenly distributed over the four randomly assigned stimulus configurations: target contralateral (blue = 1251, yellow = 1171), target ipsilateral (blue = 1164, yellow = 1136). Monkey K: target contralateral (blue = 595; yellow = 554), target ipsilateral (blue = 536; yellow = 505); monkey P: target contralateral (blue = 656; yellow = 617), target ipsilateral (blue = 628; yellow = 631). These periods were subjected to time-frequency analysis based on an epoch length of 500 ms and a step size of 50 ms.

### Spectral analysis of power and phase locking

Spectral analysis proceeded with transformation of the 500 ms epochs (as defined above) to the frequency domain via the multitaper method (MTM). We used 3 tapers, which provided a spectral smoothing of ± 4 Hz (46, 47). Epochs were zero-padded to 1 s resulting in a frequency resolution of 1 Hz. The spectral power was derived as the squared magnitude of the complex Fourier coefficients. The percentage power change from baseline was computed as:

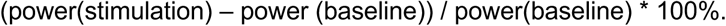

The baseline value was computed as the average value over the period from -200 ms to 0 ms before cue onset, averaged over time points and all trials from all conditions, per site.

Phase locking was quantified with the pairwise phase consistency (PPC) metric (21). PPC is not biased by the number of epochs, whereas the more conventional coherence metric has that bias. Essentially, the PPC calculation proceeds in two steps. First, the relative phases are calculated for the multiple epochs of the two signals. The second step is the crucial step: In conventional coherence calculation, those relative phases are averaged, which leads to the bias by epoch number; in PPC calculation, all possible pairs of relative phases are formed, the cosines between those relative phases are determined and those cosine values are averaged.

### Identification of spectral peaks

The PPC spectra between all site pairs were used to identify narrow-band oscillations. A phase locking metric was selected to assess oscillatory content, because it is not corrupted by 1/f background noise, and therefore provides a robust estimation of peak heights across frequency (Fig. 3A,B). Peaks were assessed using Gaussian fits to the PPC spectrum of each site pair across the ECoG grid, using the findpeaksG.m algorithm by T.C O’Haver. Each peak was assessed for statistical significance (p<0.05) via comparison to a distribution of the maximum PPC value across frequencies and site pairs to control for multiple comparisons. One-hundred random permutations of the trial order across pairs were performed followed by computation of the PPC. This procedure disrupts the phase relations across trials for each site, giving an estimate of the maximal peak height expected by chance. The probability of a peak at each frequency was found via computation of the smoothed peak histogram (Fig. 3A,B, upper panels) at each frequency and identifying the dominant peaks using the findpeaksG.m algorithm. Frequencies of interest (FOIs) were then defined as the full-width-at-half-maximum of the estimated center frequency of each peak. This revealed four peaks in each monkey, namely theta (monkey K: 2:4.4:7 Hz [start:center:end], monkey P: 2:4.8:7 Hz), beta (K: 16:18.1:20 Hz, P: 11:13.9:16 Hz), high-beta (K: 31:33.9:36 Hz, P: 29:31.6:35 Hz), and gamma (K: 73:74.5:76 Hz, P: 60:63.0:66 Hz).

### Statistical inference on power-change time courses

Statistical comparisons of time-resolved power differences were computed via permutation statistics. This entailed randomly assigning each trial to one of the four unique stimulus conditions shown in Figure 2E, while maintaining the sample sizes for each condition. Spectral analysis was then performed as described followed by the computation of each contrast (Figure 2E). This procedure was repeated 10000 times to produce a randomization distribution for both contrasts. To control for multiple comparisons, a hybrid method was employed that controls for the temporal, frequency and spatial dimensions. The multiple comparisons across the temporal and spatial dimensions, were controlled for by a max-based method (48). The multiple comparisons across the frequency dimension was controlled for by Bonferroni correction, because this is less affected by large differences in effect size across the different FOIs. Thus, the two-tailed significance criterion of 0.05, 0.025 per tail, was divided by 4 to account for the 4 FOIs tested. This resulted in the following procedure: 1) Randomization distributions were computed for each contrast, then averaged over monkeys, 2) The maximum absolute value of the power difference contrasts was found for each FOI across time windows and brain areas. 3) A critical value was then selected for each FOI from these distributions as the 99.38^th^ percentile, derived as a 4-fold correction of the two-tailed 0.05 p-value, 4) The observed power differences between contrasts, averaged across monkeys were then compared to their respective distributions to assess statistical significance at a level of p = 0.05 two-tailed.

## Author contributions

Conceptualization: C.G.R., C.A.B., and P.F.; Methodology: C.G.R., C.A.B., and P.F.; Software: C.G.R., J.V., and J.M.S.; Investigation: C.A.B. and P.F.; Formal Analysis: C.G.R., J.V., and J.M.S.; Writing – Original Draft: C.G.R. and P.F.; Writing – Review & Editing: C.G.R., C.A.B., J.V., J.M.S., and P.F.; Supervision: P.F.; Funding Acquisition: C.A.B. and P.F.

## Acknowledgements

PF acknowledges funding by DFG (SPP 1665, FOR 1847, FR2557/5-1-CORNET, FR2557/6-1-NeuroTMR), EU (HEALTH-F2-2008-200728-BrainSynch, FP7-604102-HBP, FP7-600730-Magnetrodes), a European Young Investigator Award, National Institutes of Health (1U54MH091657-WU-Minn-Consortium-HCP), and the LOEWE program (NeFF). CR acknowledges support by the Basque Government through the BERC 2018-2021 program and by the Spanish State Research Agency through BCBL Severo Ochoa excellence accreditation SEV-2015-0490. CAB acknowledges support by the FLAG-ERA JTC 2015 project CANON (co-funded by NWO). JMS acknowledges funding by NWO (VIDI-grant, 864.14.011).

## Reference

1. Yantis S, et al. (2002) Transient neural activity in human parietal cortex during spatial attention shifts. Nat Neurosci 5(10):995–1002.

2. Grent-’t-Jong T, Woldorff MG (2007) Timing and sequence of brain activity in top-down control of visual-spatial attention. PLoS Biol 5(1):e12.

3. Müller MM, Teder-Sälejärvi W, Hillyard SA (1998) The time course of cortical facilitation during cued shifts of spatial attention. Nat Neurosci 1(7):631–634.

4. Fries P (2001) Modulation of Oscillatory Neuronal Synchronization by Selective Visual Attention. Science 291(5508):1560–1563.

5. Taylor K, Mandon S, Freiwald WA, Kreiter AK (2005) Coherent oscillatory activity in monkey area v4 predicts successful allocation of attention. Cereb Cortex 15(9):1424–1437.

6. Gregoriou GG, Gotts SJ, Zhou H, Desimone R (2009) High-frequency, long-range coupling between prefrontal and visual cortex during attention. Science 324(5931):1207–1210.

7. Bosman CA, et al. (2012) Attentional Stimulus Selection through Selective Synchronization between Monkey Visual Areas. Neuron 75(5):875–888.

8. Grothe I, Neitzel SD, Mandon S, Kreiter AK (2012) Switching neuronal inputs by differential modulations of gamma-band phase-coherence. 32(46):16172–16180.

9. Siegel M, Donner TH, Oostenveld R, Fries P, Engel AK (2008) Neuronal synchronization along the dorsal visual pathway reflects the focus of spatial attention. Neuron 60(4):709–719.

10. Womelsdorf T, Fries P, Mitra PP, Desimone R (2006) Gamma-band synchronization in visual cortex predicts speed of change detection. Nature 439(7077):733–736.

11. Bichot NP, Rossi AF, Desimone R (2005) Parallel and serial neural mechanisms for visual search in macaque area V4. Science 308(5721):529–534.

12. Buffalo EA, Fries P, Landman R, Buschman TJ, Desimone R (2011) Laminar differences in gamma and alpha coherence in the ventral stream. Proc Natl Acad Sci U S A 108(27):11262–11267.

13. Gregoriou GG, Gotts SJ, Desimone R (2012) Cell-type-specific synchronization of neural activity in FEF with V4 during attention. Neuron 73(3):581–594.

14. Zhou H, Schafer RJ, Desimone R (2016) Pulvinar-Cortex Interactions in Vision and Attention. Neuron 89(1):209–220.

15. Buschman TJ, Miller EK (2007) Top-down versus bottom-up control of attention in the prefrontal and posterior parietal cortices. Science 315(5820):1860–1862.

16. Bastos AM, et al. (2015) Visual areas exert feedforward and feedback influences through distinct frequency channels. Neuron 85(2):390–401.

17. Richter CG, Thompson WH, Bosman CA, Fries P (2017) Top-Down Beta Enhances Bottom-Up Gamma. J Neurosci 37(28):6698–6711.

18. Spyropoulos G, Bosman CA, Fries P (2018) A theta rhythm in macaque visual cortex and its attentional modulation. Proc Natl Acad Sci U S A:201719433.

19. Vinck M, Womelsdorf T, Buffalo EA, Desimone R, Fries P (2013) Attentional modulation of cell-class-specific gamma-band synchronization in awake monkey area v4. Neuron 80(4):1077–1089.

20. Pesaran B, et al. (2018) Investigating large-scale brain dynamics using field potential recordings: analysis and interpretation. Nat Neurosci 21(7):903–919.

21. Vinck M, van Wingerden M, Womelsdorf T, Fries P, Pennartz CMA (2010) The pairwise phase consistency: a bias-free measure of rhythmic neuronal synchronization. Neuroimage 51(1):112–122.

22. Buffalo EA, Fries P, Landman R, Liang H, Desimone R (2010) A backward progression of attentional effects in the ventral stream. Proc Natl Acad Sci U S A 107(1):361–365.

23. Engel AK, Fries P (2010) Beta-band oscillations-signalling the status quo? Curr Opin Neurobiol. doi:10.1016/j.conb.2010.02.015.

24. Landau AN, Fries P (2012) Attention Samples Stimuli Rhythmically. Curr Biol 22(11):1000–1004.

25. Rollenhagen JE, Olson CR (2005) Low-Frequency Oscillations Arising From Competitive Interactions Between Visual Stimuli in Macaque Inferotemporal Cortex. J Neurophysiol 94(5):3368–3387.

26. Ghose GM, Ts’o DY (2017) Integration of color, orientation, and size functional domains in the ventral pathway. Neurophotonics 4(3):031216.

27. Kotake Y, Morimoto H, Okazaki Y, Fujita I, Tamura H (2009) Organization of color-selective neurons in macaque visual area V4. J Neurophysiol 102(1):15–27.

28. Logothetis NK, Pauls J, Augath M, Trinath T, Oeltermann A (2001) Neurophysiological investigation of the basis of the fMRI signal. Nature 412(6843):150–157.

29. Niessing J, et al. (2005) Hemodynamic signals correlate tightly with synchronized gamma oscillations. Science 309(5736):948–951.

30. Scheeringa R, et al. (2011) Neuronal dynamics underlying high- and low-frequency EEG oscillations contribute independently to the human BOLD signal. Neuron 69(3):572–583.

31. Scheeringa R, Fries P (2019) Cortical layers, rhythms and BOLD signals. Neuroimage 197:689–698.

32. Pesaran B, Pezaris JS, Sahani M, Mitra PP, Andersen RA (2002) Temporal structure in neuronal activity during working memory in macaque parietal cortex. Nat Neurosci 5(8):805–811.

33. Rubehn B, Bosman C, Oostenveld R, Fries P, Stieglitz T (2009) A MEMS-based flexible multichannel ECoG-electrode array. J Neural Eng 6(3):036003.

34. Pinotsis DA, et al. (2014) Contrast gain control and horizontal interactions in V1: A DCM study. Neuroimage 92:143–155.

35. Brunet N, Vinck M, Bosman CA, Singer W, Fries P (2014) Gamma or no gamma, that is the question. Trends Cogn Sci (Regul Ed) 18(10):507–509.

36. Brunet N, et al. (2014) Stimulus repetition modulates gamma-band synchronization in primate visual cortex. Proc Natl Acad Sci USA 111(9):3626–3631.

37. Brunet N, et al. (2015) Visual Cortical Gamma-Band Activity During Free Viewing of Natural Images. Cereb Cortex 25(4):918–926.

38. Richter CG, Thompson WH, Bosman CA, Fries P (2015) A jackknife approach to quantifying single-trial correlation between covariance-based metrics undefined on a single-trial basis. Neuroimage 114:57–70.

39. Vinck M, et al. (2015) How to detect the Granger-causal flow direction in the presence of additive noise? Neuroimage 108:301–318.

40. Bastos AM, et al. (2015) A DCM study of spectral asymmetries in feedforward and feedback connections between visual areas V1 and V4 in the monkey. Neuroimage 108:460–475.

41. Lewis CM, Bosman CA, Womelsdorf T, Fries P (2016) Stimulus-induced visual cortical networks are recapitulated by spontaneous local and interareal synchronization. Proc Natl Acad Sci USA 113(5):E606–15.

42. Oostenveld R, Fries P, Maris E, Schoffelen J-M (2011) FieldTrip: Open source software for advanced analysis of MEG, EEG, and invasive electrophysiological data. Comput Intell Neurosci 2011:156869.

43. Van Essen DC (2012) Cortical cartography and Caret software. Neuroimage 62(2):757–764.

44. Rohlfing T, et al. (2012) The INIA19 Template and NeuroMaps Atlas for Primate Brain Image Parcellation and Spatial Normalization. Front Neuroinform 6:27.

45. Welch P (1967) The use of fast Fourier transform for the estimation of power spectra: A method based on time averaging over short, modified periodograms. IEEE Trans Audio Electroacoust 15(2):70–73.

46. Thomson DJ (1977) Spectrum Estimation Techniques for Characterization and Development of WT4 Waveguide-I. Bell System Technical Journal 56(9):1769–1815.

47. Percival DB, Walden AT (1993) Spectral Analysis for Physical Applications (Cambridge University Press).

48. Nichols TE, Holmes AP (2002) Nonparametric permutation tests for functional neuroimaging: a primer with examples. Hum Brain Mapp 15(1):1–25.

